# Study on Spatio-temporal Prevalence and Hematological Attributes of Bovine Babesiosis in Cattle Population of Layyah, Southern Punjab, Pakistan

**DOI:** 10.1101/2023.01.09.523272

**Authors:** Absar Ahmad, Zulfiqar Ali, Mushtaq Hussain Lashari

## Abstract

The objective of the present study was to investigate the spatiotemporal prevalence of bovine babesiosis in cattle population (n=376) of District Layyah, South Punjab, Pakistan as affected by location, age, breed, gender and seasons. Blood samples were collected aseptically and assessed for babesiosis through microscopy and PCR, and through automated analyzer for hematological attributes. Overall results of prevalence as attained through PCR in cattle population showed a significantly (P≤0.05) lower prevalence of 19.4% (n=72/376) as compared to 37.2% (n=140/276) through microscopy. None of the studied cattle from Cholistani breed were Babesia-positive. However, significantly (P≤0.05) higher prevalence was noticed for crossbred cattle (46.7%, n= 50/107) followed by that in Friesian (16.1%, n= 10/62), Jersey (7%, n= 5/71) and Sahiwal (6.9%, n= 7/101) cattle breeds. Female cattle (19.5%, n= 55/281) and age group 1 (Up to 2 years) (40%, n= 42/105) had higher prevalence of Babesia as ascertained through PCR in comparison to their counterpart groups. Significantly (P≤0.05) higher prevalence of 35.9% (n=60/167) was shown in summer as compared to that in winter season (5.7%, n= 12/209). All the positive samples produced the 490bp amplicons specific and typical for Babesia bigemina. Regarding hematology, hemoglobin concentration, erythrocytic count, hematocrit and mean corpuscular volume were significantly (P≤0.05) lower in babesia-positive cattle as compared to healthy ones. In a nutshell, indigenous cattle breeds are tick-resistant hardy breeds and do not show severe signs of babesiosis as compared to exotic and crossbred cattle. Furthermore, Southern Punjab area of Pakistan has a different spatiotemporal distribution of babesiosis with bigemina being predominant.

## Introduction

Bovine babesiosis is highly pathogenic and the signs and symptoms of its infection are varied depending upon the infected species, age, climatic conditions of the area, tick population, and immune system of the host. However, the general indications include fever, anemia, jaundice and hemoglobinurea. Though the signs and symptoms of its infection are varied depending upon the infected species, age, climatic conditions of the area, tick population, and immune system of the host as mentioned above, however, various techniques (both conventional and molecular) are in vogue. Traditionally, its diagnosis is attained through direct identification of the parasite in infected RBCs of the host, using appropriately-stained blood smears. Infected RBCs are hard to find in acute infections owing to sequestration of RBCs to the walls of the capillaries, hence serological (Immunoblotting and immunofluorescent testing) and molecular diagnostic tests (standard PCR, nested PCR, rtPCR, reverse line blot, loop-mediated isothermal amplification) have made a stronger footing being highly specific, sensitive and precise (Ozubek et al., 2020; Sivakumar et al., 2020).

Over the years, livestock has emerged as a main sub-sector of agriculture in Pakistan with its share of 60.09% to agriculture value addition and 11.5% to the GDP during the financial year 2021 (Finance Division, 2021). A growth of 3.06% in this sub-sector has been reported by the Government of Pakistan. Livestock engages more than 8 million rural families providing them with 35-40% of their income. Hence, it can be dubbed as the most vital sector for the socio-economic uplift of rural communities in specific and whole country in general.

The geographical location of Pakistan being in the Warm Climate Zones (WCZs) of the world as per UNO is endemic to high prevalence of tick-borne diseases especially babesiosis. Lowered productivity and high mortality as a feature of this infection imply a grave economic burden especially on rural livestock farmers of the country. Extensive studies have been conducted regarding tick prevalence in Pakistan with highest prevalence reported for cattle (28.2%) by that in sheep (18.8%), buffaloes (14.7%) and goats (12.3%) (Ghosh et al., 2007; Jabbar et al., 2015). These varying levels of tick prevalence in Pakistan have been attributed to varying geoclimatic conditions, lifestyles of livestock keepers, livestock production systems, farmer awareness, management practices and governmental interventions (Karim et al., 2017; Muhammad, Naureen, Firyal, & Saqib, 2008). The warm climate of the country allied with higher tick prevalence makes it vulnerable to bovine babesiosis. Extensive research work has been reported on prevalence of bovine babesiosis from various cities and provinces of Pakistan. From Punjab, Pakistan, the research work on its prevalence has mostly emanated from cities of upper Punjab. However, not much work has yet been carried out in cities of Lower/Southern Punjab (a newly proposed province of Pakistan) which includes 13 major cities. The present research work has thus been devised with an aim to assess situation analysis/spatiotemporal prevalence of bovine babesiosis using both traditional diagnostic technique and PCR in various cattle breeds (Sahiwal, Cholistani, Crossbred, Friesian, Jersey) being reared in district Layyah of Southern Punjab, Pakistan. Furthermore, it also aims to compare hematological attributes between Babesia-infected and non-infected animals.

## Materials and Methods

### Geo-location of the Study

The study was conducted at the District Layyah, Southern Punjab, Pakistan. Layyah is 72^nd^ biggest city of Pakistan lying within the Dera-Ghazi-Khan Division (Figure 1). Covering an area of 6291 km^2^, it lays at latitude of 30° N and longitude of 70° E. Its geographic characteristic is sandy land as it is entrapped between River Indus and River Chenab. It has a dry climate with scanty rainfall with a wide temperature variation of 7-41°C with an average temperature of 25.2°C (S. Ashraf et al., 2021). In summers the temperature may reach 48°C. The recorded annual precipitation for Layyah is 236.3mm.

**Figure 1:**
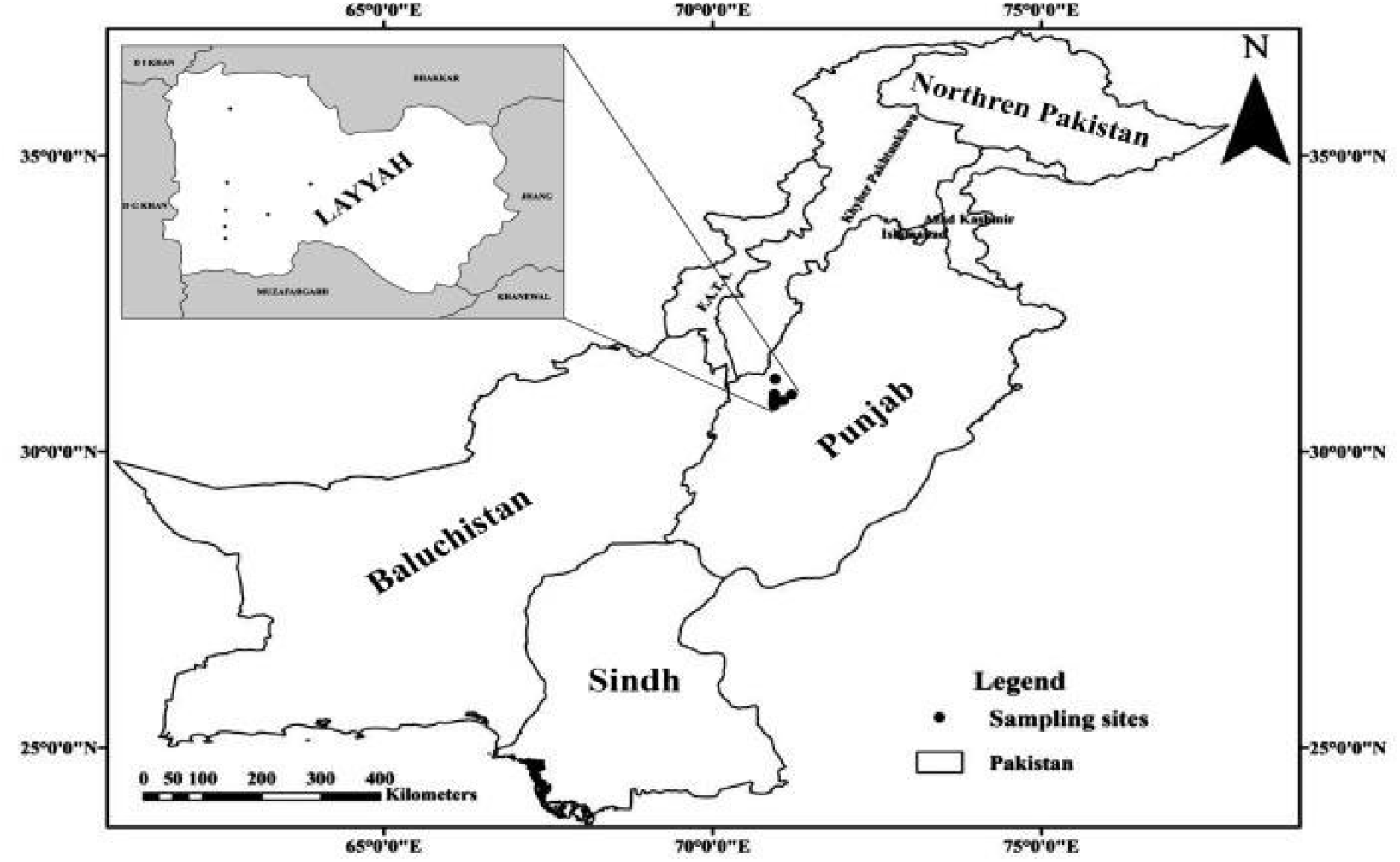
Geo-location of Layyah District, Southern Punjab, Pakistan.

### Study Animals and Grouping

District Layyah has three Tehsils namely Layyah, Karor and Choubara and 44 Union Councils (Figure 2). Private livestock farms were identified and registered under the study from each of the three Tehsils. Qualitative/quantitative data regarding the traits of growth, performance, nutritional plain, housing, production systems, health status, veterinary services provision and identification of constraints for the bovine rearing were acquired through structural interviews with owners, predefined questionnaires, focus group discussions, and participatory rural appraisal.

**Figure 2.**
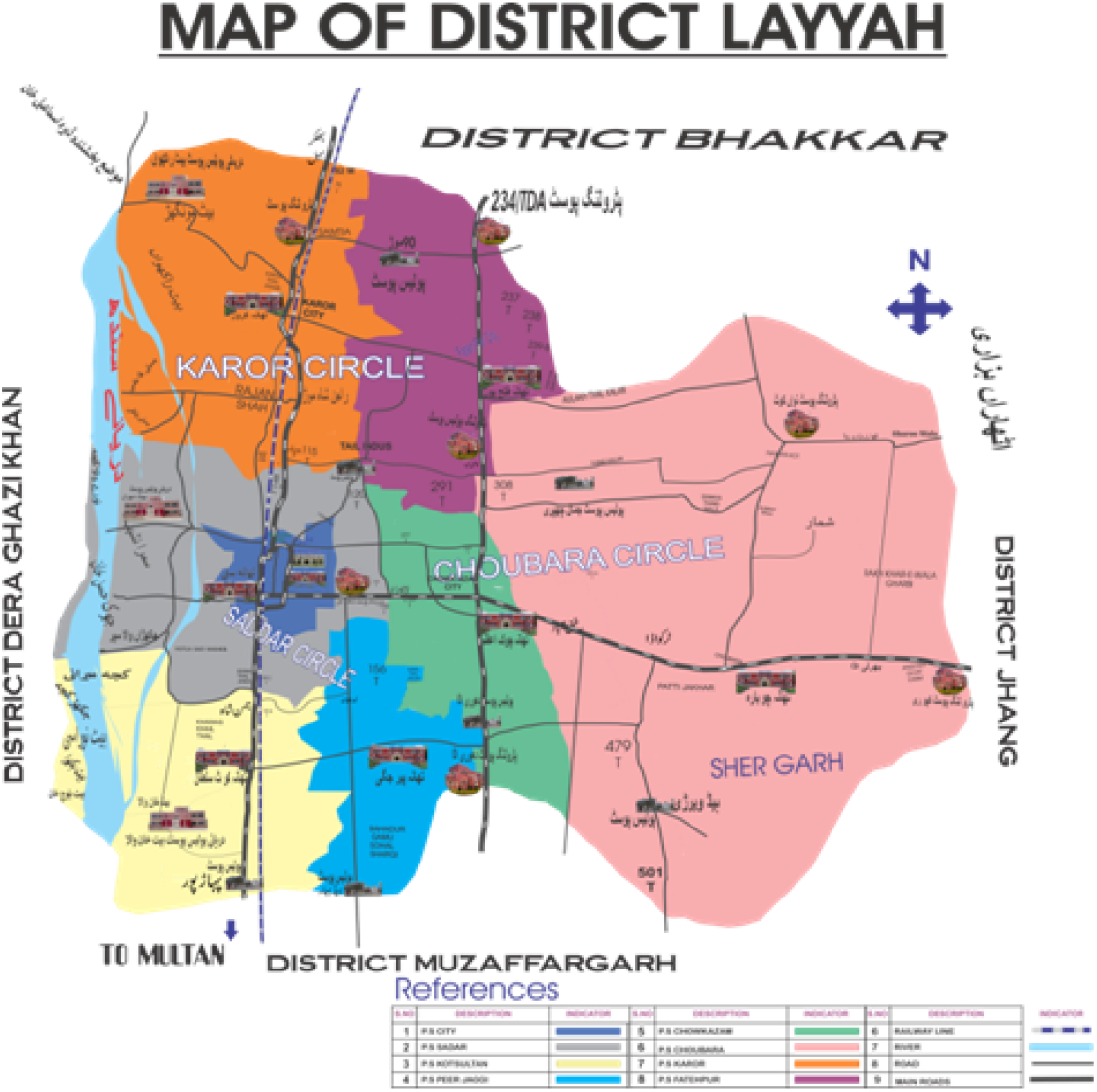
Map of Layyah indicating its three Tehsils and 44 Union Councils.

The cattle population (n=376) of both genders and belonging to different age and breed groups was randomly selected and sampled throughout the year of 2021 in order to ascertain the effect of location, age, breed, gender and seasons. The tehsil-wise study population of bovines is presented in Table 1 whereas the age, gender, breed and season-wise study population of cattle is given in Table 2.

**Table 1.**
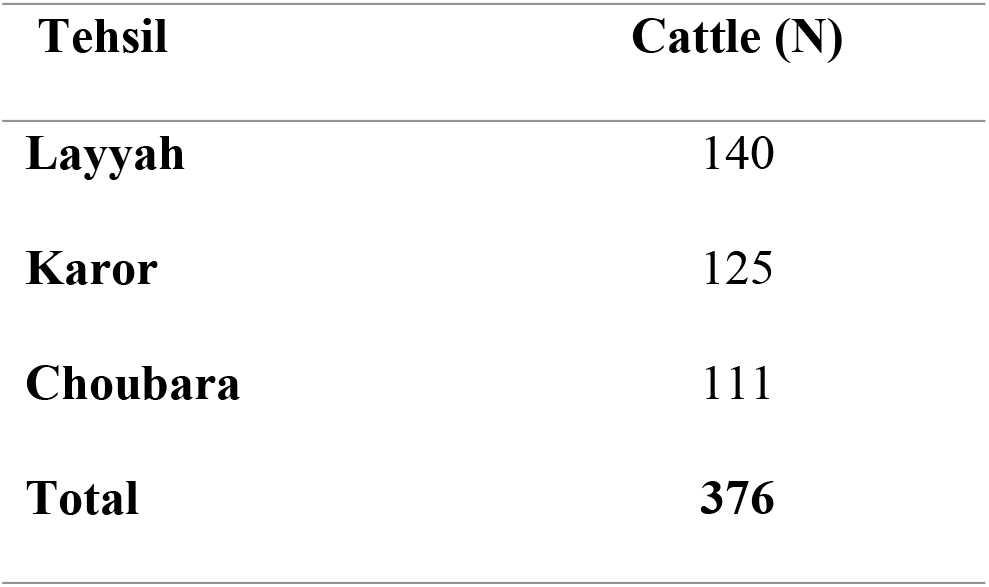
The tehsil-wise cattle population (n=376) incorporated in the study.

| **Tehsil** | **Cattle (N)** |
| --- | --- |
| **Layyah** | 140 |
| **Karor** | 125 |
| **Choubara** | 111 |
| **Total** | **376** |

**Table 2.** The age, gender and breed-wise grouping of cattle (n=376) population incorporated in the study.

| **Age** |  |  | **Gender** |  | **Breed** |  |  |  |  | **Seasons** |  |
| --- | --- | --- | --- | --- | --- | --- | --- | --- | --- | --- | --- |
| **G1** | **G2** | **G3** | **Males** | **Females** | **Sahiwal** | **Cholistani** | **Crossbred** | **Friesian** | **Jersey** | **S** | **W** |
| **105** | 152 | 119 | 95 | 281 | 101 | 35 | 107 | 62 | 71 | 167 | 209 |

G1= Up to 02 Years, G2= 03-04 Years, G3= 05-06 Years
S= Summer (April to September), W= Winter (October to March)

### Clinical Examination and Blood Collection

Prior to blood collection, a thorough on-site clinical examination was carried out on each animal and deductions were made on basis of anamnesis, clinical examination, and signs and symptoms. Prevalence of ticks on animal body was confirmed through visual and manual inspection. Blood was collected aseptically in three aliquots from the coccygeal vein of the animal after a thorough restraint and was placed in EDTA containing vacutainers (BD Vacutainers®, Becton Dickinson, USA). Timing, personnel, and method of restraint for blood collection were maintained same throughout the study period in order to minimize stress in animals (Figure 3.3). The three aliquots of each blood sample from an animal were transported in ice packs to: a) Paraveterinary Institute, Karor Lal Easan, Layyah, South Punjab, (sub-campus of University of Veterinary and Veterinary Animal Sciences, Lahore, Pakistan) for slide preparation, microscopy and hematological analysis, b) College of Veterinary and Animal Sciences (UVAS), Jhang (sub-campus of University of Veterinary and Veterinary Animal Sciences, Lahore, Pakistan) for conventional PCR.

**Figure 3.**
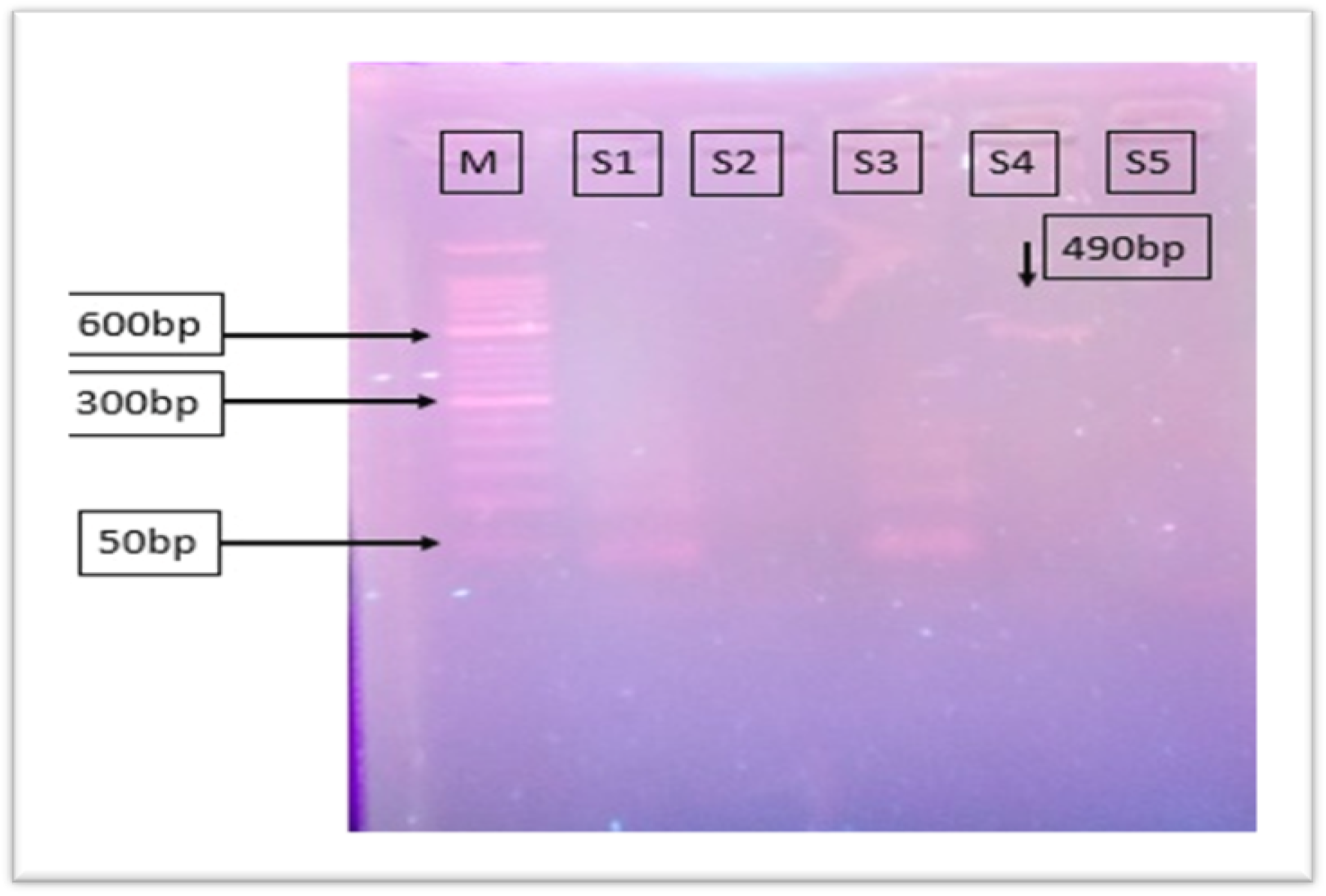
Representative gel of the samples showing specificity of 490amplicon for *Babesia bigemina* in infected animals.

### Hematological Analyses

A thin blood smear was prepared on clean, grease-free glass slides, air dried, fixed through methanol and stained using Field’s stain (Umair Riaz, 2021). This blood film was examined under the oil immersion lens of a trinocular microscope (100X) fitted with a 16 MP camera (LaboMed Inc. LB 243, USA) for presence of *babesia* species and morphological alterations of blood cells. Various hematological attributes *viz*. Hemoglobin concentration (Hb), Total Erythrocyte Count (TEC), Packed Cell Volume (PCV), Total Leukocyte Count (TLC), Neutrophils, Eosinophils, Basophils, Monocytes and Lymphocytes were analyzed through a veterinary hematology analyzer (Vet Exigo, H400, Boule Diagnostics AB, Sweden).

### Molecular Diagnostics

DNA extraction was carried out using a commercial kit (Gene Jet Whole Blood DNA Purification Kit, Thermo-fisher Scientific, USA; catalogue no. K0782) following the procedure described by the manufacturer. Briefly, 200µL of whole blood sample was added in a 1.5mL eppendorf tube along with 400µL of lysis solution and 20µL of Proteinase-K. This was incubated at 56°C for 10 minutes in shaking water-bath after vortexing the mixture. After addition of 200 µL of 96-100% ethanol, the mixture was transferred in Gene JET Genomic DNA purification column inserted in a collection tube. The column was centrifuged after addition of ethanol, wash buffer 1 and wash buffer 2 at different speed and time intervals for removal of debris. Elution buffer was added to elute the DNA from silicon sieve of spin column. Finally, the DNA was collected in 1.5 ml Eppendorf tubes after centrifugation and stored in −20 °C freezer for further processing.

For conventional PCR, BJ1 (5’-GTC-TTG-TAA-TTG-GAATGA-TGG-3’) and BN2 (5’-TAG-TTT-ATG-GTT-AGG-ACT-ACG-3’) primers were used along with PCR Master Mix (2 × Ace-Taq Master Mix (Dye Plus) P412). The PCR mixture of 20µL was comprised of (PCR master mix 10µL, Primer BJ1 1µL, Primer BN2 1µL, Template DNA 5µL, and nuclease free water 3µL). A total of 35 cycles (Initial heating and denaturation at 95°C for 3 minutes, denaturation at 95°C for 30sec, annealing at 52°C for 30sec, extension at 72°C for 30sec and final extension at 72°C for 5minutes) were conducted in Thermocycler (Bio-Rad, USA). Positive control samples were taken from Molecular Parasitology Laboratory, Veterinary Faculty, UAF, Pakistan, whereas sterile distilled water was used as a negative control.

The PCR products along with positive and negative controls were analysed using 50 bp and 100bp DNA ladder on 2% agarose gel containing ethidium bromide at the rate of 0.5µg/µL of gel in 1X TAE buffer. About 10 µL of PCR product was loaded in agarose gel for visualization. The gel electrophoresis was performed in gel documentation system (Bio-Rad, USA) at 110V, and 400 amp (maximum) for 35 minutes or until the dye migrated into two third of gel. Finally, the gel was visualized, and image was taken with the help of UV illuminator (Biostep, Germany).

### Statistical Analyses

All the collected data was sorted on the basis of location, age, breed, gender and seasons. Regarding age, the animals were grouped as G1= Up to 02 Years, G2= 03-04 Years, and G3= 05-06 Years. The cattle breeds included Sahiwal (n=101), Cholistani (n=35), Crossbred (n=107), Friesian (n=62) and Jersey (n=71). The seasons were designated as summer (April to September), and winter (October to March). Descriptive statistics was implied to attain frequencies, percentages and measures of central tendency. Regarding prevalence, percentages were deduced and results were presented as odds ratio with their 95 % confidence intervals. Difference of prevalence between location, age, breed, gender and seasons was deducted through Chi-square test keeping P≤0.05 as statistically significant. All the hematological attributes were expressed as means (±SE). The difference of hematological attributes between Babesia-positive and Babesia-negative animals was deducted through unpaired t-test. Predictive values were determined through sensitivity and specificity of blood smear examination and PCR test. All above statistical analyses were carries out through Statistical Package for Social Sciences (V19, IBM, USA).

## Results

### Clinical Observations

The on-site diagnosis/field survey revealed that regarding cattle breeds, the crossbred, Friesian and Jersey cattle showed severe and typical clinical signs of pyrexia, anorexia, nervous signs, elevated pulse rate, anemia, hyperpnea, jaundice, pale conjunctiva/vaginal membrane and hemoglobinurea. Generalized poor demeanor of the body was also observed in some animals. The Sahiwal cattle, on the other hand, revealed milder symptoms of pyrexia and anorexia only. On the contrary, it was noticed that none of the Cholistani breed of cattle showed any of the typical signs and symptoms related to babesiosis.

### Prevalence through Stained Blood Smear Microscopy

An overall prevalence of 37.2% (n=140/376) was observed in cattle population of the present study using stained blood smear microscopy. Breed-wise results for cattle indicated an overall higher (P≤0.05) prevalence of 64.5% (n=44/62) for Friesian followed by 51.4% (n=55/107), 43.6% (n=31/71) and 7.9% (n=8/101) for crossbred, Jersey and Sahiwal breeds of cattle, respectively. Lowest prevalence was noticed for Cholistani cattle breed bring 5.7% (n=2/35). Regarding age, Group 1 (Up to 02 Years) had significantly (P≤0.05) higher prevalence of 57.1% (n=60/105) followed by 39.4% (n=60/152) and 16.8% (n=20/119) for Group 2 and 3, respectively. Females had significantly (P≤0.05) higher prevalence (42.7%, n=120/281) as compared to that in males (21.0%, n=20/95). Significantly (P≤0.05) higher prevalence of bovine babesiosis in cattle was noticed during summer season being 47.9% (n=80/167) as compared to 28.7% (n=60/209) in winter season.

### Prevalence through PCR

Overall results of prevalence as attained through PCR in cattle population (n=376) showed a significantly (P≤0.05) higher prevalence of 19.4% (n=72/376) through PCR as compared to 37.2% (n=140/276) through microscopy. Breed, tehsils, gender, seasons and age-wise results have been tabulated in Table 3. None of the studied cattle from Cholistani breed were Babesia-positive. However, significantly (P≤0.05) higher prevalence was noticed for crossbred cattle (46.7%, n= 50/107) followed by that in Friesian (16.1%, n= 10/62), Jersey (7%, n= 5/71) and Sahiwal (6.9%, n= 7/101) cattle breeds. Regarding tehsils, Layyah had highest prevalence of 30.7% (n=43/140) followed by the other two tehsils of Karor (14.4%, n= 18/125) and Choubara (9.9%, n= 11/111). Female cattle (19.5%, n= 55/281) and age group 1 (Up to 2 years) (40%, n= 42/105) had higher prevalence of Babesia as ascertained through PCR in comparison to their counterpart groups. Similarly, significantly (P≤0.05) higher prevalence of 35.9% (n=60/167) was shown in summer as compared to that in winter season (5.7%, n= 12/209).

**Table 3.**
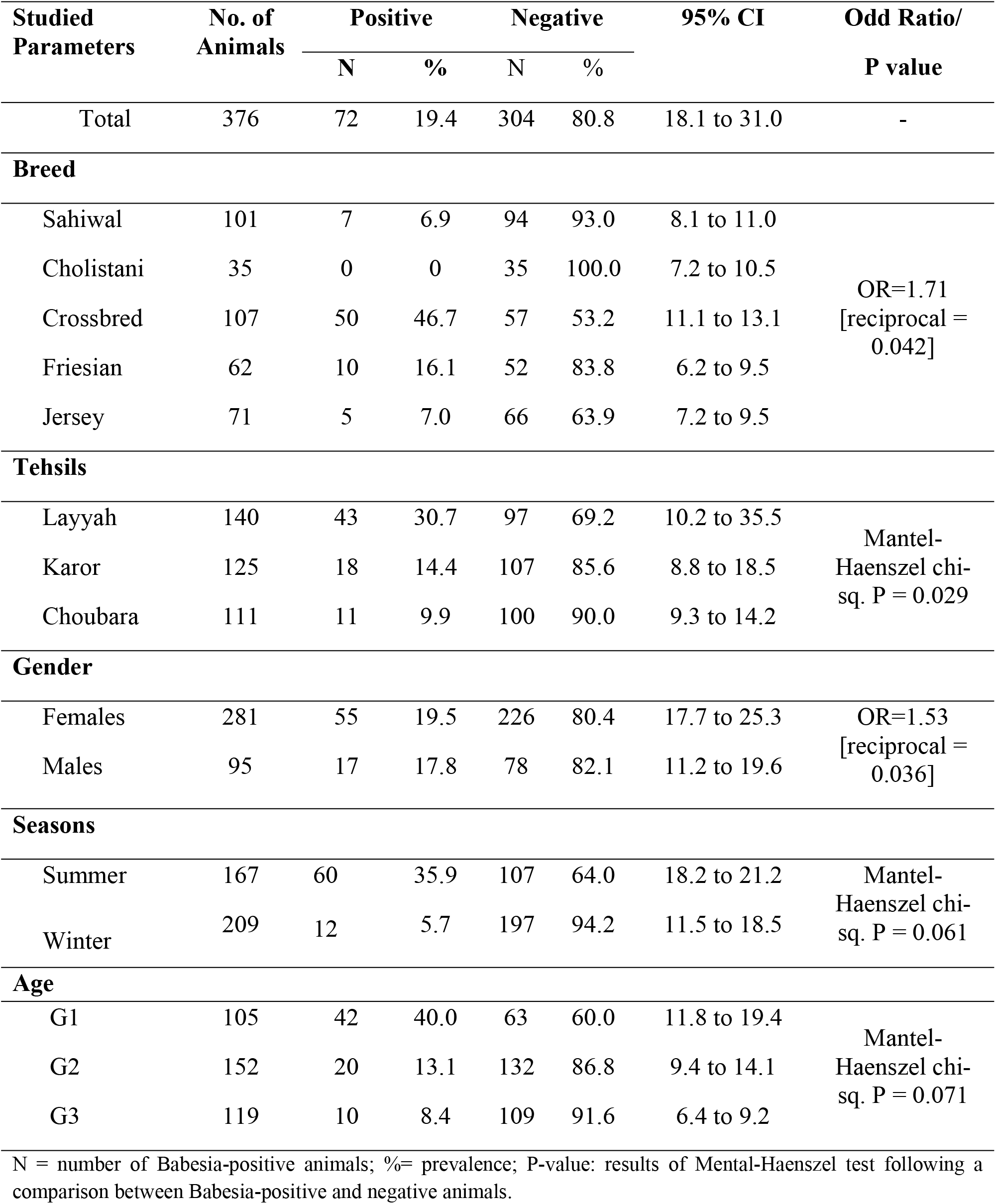
Overall prevalence of bovine babesiosis in cattle population (n=376) for Layyah District, Southern Punjab, Pakistan as ascertained through PCR.

| Studied Parameters | No. of Animals | Positive |  | Negative |  | 95% CI | Odd Ratio/<br>P value |
| --- | --- | --- | --- | --- | --- | --- | --- |
|  |  | N | % | N | % |  |  |
| Total | 376 | 72 | 19.4 | 304 | 80.8 | 18.1 to 31.0 | - |
| Breed |  |  |  |  |  |  |  |
| Sahiwal | 101 | 7 | 6.9 | 94 | 93.0 | 8.1 to 11.0 | OR=1.71<br>[reciprocal = 0.042] |
| Cholistani | 35 | 0 | 0 | 35 | 100.0 | 7.2 to 10.5 |  |
| Crossbred | 107 | 50 | 46.7 | 57 | 53.2 | 11.1 to 13.1 |  |
| Friesian | 62 | 10 | 16.1 | 52 | 83.8 | 6.2 to 9.5 |  |
| Jersey | 71 | 5 | 7.0 | 66 | 63.9 | 7.2 to 9.5 |  |
| Tehsils |  |  |  |  |  |  |  |
| Layyah | 140 | 43 | 30.7 | 97 | 69.2 | 10.2 to 35.5 | Mantel-Haenszel chi-sq. P = 0.029 |
| Karor | 125 | 18 | 14.4 | 107 | 85.6 | 8.8 to 18.5 |  |
| Choubara | 111 | 11 | 9.9 | 100 | 90.0 | 9.3 to 14.2 |  |
| Gender |  |  |  |  |  |  |  |
| Females | 281 | 55 | 19.5 | 226 | 80.4 | 17.7 to 25.3 | OR=1.53<br>[reciprocal = 0.036] |
| Males | 95 | 17 | 17.8 | 78 | 82.1 | 11.2 to 19.6 |  |
| Seasons |  |  |  |  |  |  |  |
| Summer | 167 | 60 | 35.9 | 107 | 64.0 | 18.2 to 21.2 | Mantel-Haenszel chi-sq. P = 0.061 |
| Winter | 209 | 12 | 5.7 | 197 | 94.2 | 11.5 to 18.5 |  |
| Age |  |  |  |  |  |  |  |
| G1 | 105 | 42 | 40.0 | 63 | 60.0 | 11.8 to 19.4 | Mantel-Haenszel chi-sq. P = 0.071 |
| G2 | 152 | 20 | 13.1 | 132 | 86.8 | 9.4 to 14.1 |  |
| G3 | 119 | 10 | 8.4 | 109 | 91.6 | 6.4 to 9.2 |  |

N = number of Babesia-positive animals; %= prevalence; P-value: results of Mental-Haenszel test following a comparison between Babesia-positive and negative animals.

**Table 4.** Overall mean (±SE) values of hematological attributes between Babesia-positive and negative cattle of Layyah District, Southern Punjab, Pakistan.

| Attributes | Positive<br>(n= 72) | Negative<br>(n= 304) | Overall<br>(n= 376) |
| --- | --- | --- | --- |
| Hemoglobin (g/dL) | 9.6±0.7 | 12.8±0.9* | 13.2±0.4 |
| Total erythrocyte count ($10^{12}$/L) | 6.9±0.4 | 8.8±0.4* | 7.2±0.7 |
| Packed cell volume (%) | 32.0±1.1 | 39.9±1.8* | 37.2±1.7 |
| Mean corpuscular volume (fL) | 48±2.1 | 55±1.2* | 48.2±1.4 |
| Mean corpuscular hemoglobin (pg) | 21.2±0.8 | 20.8±0.5 | 21.2±0.7 |
| Mean corpuscular hemoglobin concentration (g/L) | 330.7±10.2 | 329.1±11.5 | 320.7±10.2 |
| Total leukocyte count ($10^9$/L) | 11.4±0.4 | 9.2±0.4* | 11.2±0.9 |
| Neutrophils (%) | 44.5±0.9 | 42.2±1.1 | 43.2±1.7 |
| Lymphocytes (%) | 58.7±1.9 | 41.8±0.9* | 52.7±2.7 |
| Eosinophils (%) | 2.3±0.7 | 1.5±0.4 | 1.9±0.01 |
| Basophils (%) | 2.9±0.7 | 2.7±0.4 | 1.9±0.07 |
| Monocytes (%) | 1.4±0.1 | 1.1±0.1 | 1.1±0.04 |

\*P ≤ 0.05 within Babesia-positive and negative animals within the rows

### Molecular Diagnostics

The results of prevalence attained through stained slide microscopy and PCR were significantly different (P≤0.05) from each other being higher for microscopy. All the positive samples produced the 490bp amplicons specific and typical for *Babesia bigemina* (Figure 3).

The sensitivity, specificity, positive predictive and negative predictive values for blood smear examination were 8.9, 45.1, 1.8 and 80.6%, respectively. Similar values for PCR were 91.0, 54.8, 19.3 and 98.1%, respectively.

### Hematological Profile

The overall mean (±SE) values of various hematological attributes in Babesia-positive and negative cattle are given in Table. All the values of healthy non-infected cattle were within the reference ranges described previously in different studies.

Results revealed that Hb concentration, TEC, PCV and MCV were significantly (P±0.05) lower in babesia-positive cattle as compared to healthy ones. However, the TLC and lymphocytes were higher (P≤0.05) for infected cattle as compared to healthy, negative ones.

## Discussion

### Clinical Examination

Results regarding clinical examination of cattle during the on-site/field survey revealed that the crossbred, Friesian and Jersey cattle showed severe and typical clinical signs of pyrexia, anorexia, nervous disturbance, elevated pulse rate, anemia, hyperpnea, jaundice, pale conjunctiva/vaginal membrane and hemoglobinurea. Signs and symptoms shown in Babesia-infected-animals have been studied extensively and two main factors are associated with their severity being virulence of species/strain, and host vulnerability (depending upon its age, gender, season, physiological status and immunological condition) (Chaudhry, Suleman, Younus, & Aslim, 2010). The *B. bovis* is highly pathogenic and virulent as compared to *B. bigemina* as provided by earlier studies. Similarly, African, Asian and Israeli strains have been reported to be highly virulent as compared to Australian strains (Friedhoff, 2018). The signs and symptoms noticed in infected cattle of the present study coincide with earlier studies. It has also been reported that anemia allied with hemoglobinurea may cause abortions in female pregnant cattle (Iseki et al., 2010). The higher severity of signs and symptoms noticed in infected crossbred and exotic breeds (Friesian and jersey) of the present study are also in line with previous studies. While studying exotic cattle in Australia similar signs for babesiosis have been reported (Aziz et al., 2014). In another study it has been reported that vaccinated cattle are highly unlikely to show any symptoms of babesiosis (Schlögl et al., 2020), however it was still endorsed that all animals may thoroughly be inspected for clinical signs at the beginning of and in middle of summer season (Holzheu, Delbeccº, Baumgartner, & Hofer, 2016). A study from Pakistan, though conducted on theileriosis has also reported similar signs and symptoms for crossbred and exotic cattle breeds (SA, RS, & MM, 2016). Similar study conducted on prevalence of babesiosis in Friesian and Jersey cattle breeds of Livestock Experimental Station, Bhunike, Punjab, Pakistan has also reported severe signs and symptoms of babesiosis in these exotic cattle breeds (Zahid, Latif, & Baloch, 2005). The severity of signs and symptoms in cross bred and exotic cattle breeds seems to be an innate genetic characteristic of these breeds. The comparative hematological profile assessed between *Bos indicus* and *Bos taurus* (exotic) cattle breeds in various studies clearly indicate substantially lower hematological attributes (especially TEC and PCV) as compared to indigenous native cattle breeds which may be a plausible cause of the severity in signs and symptoms for infected exotic cattle breeds (SA et al., 2016). This aspect of hematological differences will be discussed in detail ahead.

In the present study, the Sahiwal cattle showed milder symptoms of pyrexia and anorexia only, whereas Cholistani breed of cattle showed none of the typical signs and symptoms for babesiosis. Sahiwal and Cholistani breed of cattle are amongst the 15 indigenous, humped, zebu breeds of cattle in Pakistan (M. S. Khan, Rehman, Khan, & Ahmad, 2008). Cholistani breed of cattle is an indigenous, tick-resistant breed of cattle being reared under pastoralism in the Cholistan desert of Pakistan. A study on theileriosis conducted on this Cholistani breed of cattle (n= 264) has clearly reported that this breed is a tick-resistant and hence Babesia-resistant breed of Pakistan (SA et al., 2016). This study envisaged that a prevalence of 19.3% for noticed for theileriosis, yet none of the Cholistani cattle showed any clinical signs of theileria apart from the fact that they were tick-ridden. This character was plausibly justified with an innate ability of this breed to be tick-resistant as reported for other zebu cattle breeds throughout the world (Godfrey & Hansen, 1994; U. Farooq & Idris, 2017).

### Diagnostic Aspects

The percentage prevalence attained through PCR in the present study was 19.4% (n=72/376) which was lower than that of 37.2% (n=140/376) attained through microscopy. Furthermore, in the present study, the sensitivity, specificity, positive predictive, and negative predictive values for blood smear examination were lower than that for PCR. The traditional and conventional diagnostic test being utilized globally for diagnosis of various intra-erythrocytic diseases in bovines is blood smear microscopy. It is a routine test which needs less expertise, is cheap, and can be conducted as an infield/cow-side test. However, Babesia (and other intra-erythrocytic parasites) parasite is difficult to be viewed through microscopy if a chronic condition occurs as reported in previous studies (Chaudhry et al., 2010). Owing to a vast expansion and validation of various molecular diagnostic techniques, various types of PCRs such as nested PCR, real-time PCR, quantitative PCR and reverse-transcriptase PCR are now being used for confirmed diagnosis. Various research studies have reported higher sensitivity, specificity and repeatability for these molecular techniques as compared to blood slide microscopy (Bal, Mahajan, Filia, Kaur, & Singh, 2016; Singh, Haque, Singh, & Rath, 2013; Terkawi et al., 2011).

In our results, prevalence attained for babesiosis through blood smear microscopy was lower as compared to that attained through PCR. This is in contrast to a research work reported from Pakistan on buffaloes which reports a higher prevalence of 29% through PCR as compared to 18% for blood smear microscopy (Chaudhry et al., 2010). However, the results of sensitivity and specificity of both tests are same as those seen in our present study. Results similar to ours have also been reported in another work reported from KP province of Pakistan on cattle blood which revealed a higher sensitivity and specificity of PCR as compared to light microscopy of the blood (Pakistan, 2013). While studying theileriosis in Cholistani breed of cattle in an earlier study, sensitivity, specificity, positive predictive, and negative predictive values for blood smear examination have been reported as 8.9, 45.1, 1.8, and 80.6%, respectively. Coinciding values for PCR were 91.0, 54.8, 19.3, and 98.1%, respectively in this work. These results are also in line with our results. A study conducted in Mexico to assess comparative efficacy of IFAT, ELISA and ICT as diagnostic techniques for babesiosis has reported highest sensitivity and specificity for ICT (Lira-Amaya et al., 2021). Apart from being cheapest and fastest method of diagnosis, blood smear microscopy has lower sensitivity and hence increased chances of false positives (Mosqueda, Olvera-Ramirez, Aguilar-Tipacamu, & J Canto, 2012). From Pakistan, mostly the diagnosis of intra-erythrocytic infections such as babesiosis is mostly being carried out through blood smear microscopy with gradual incorporation of PCR (Rafique et al., 2015).

Considering the molecular diagnostics of Babesia of the present study, it was revealed that all the positive samples produced the 490bp amplicons specific and typical for *Babesia bigemina*. The present results are not in line with studies conducted in other regions/provinces of Pakistan which during molecular detection of Babesia have reported a higher incidence of *Babesia bovis* with 907bp amplicons specific to *B. bovis* (Farooqi et al., 2017; Zulfiqar et al., 2012).

### Spatiotemporal Prevalence

The overall prevalence of bovine babesiosis attained through blood smear microscopy (37.2%) was higher than that ascertained through PCR (19.4%) in the present study for cattle population (n=376) of Layyah District, South Punjab, Pakistan. A lot of work has been conducted on prevalence of babesiosis in Pakistan using both blood smear microscopy and PCR. However, none of it has emanated from Southern Punjab region of Pakistan. Comparing these results with research work conducted earlier in Pakistan, it was noticed that in contrast to our results, past research has reported higher values of prevalence through PCR as compared to those attained through blood smear microscopy (Chaudhry et al., 2010; Siddique, Sajid, Iqbal, & Saqib, 2020). A study conducted on three agro-pastoral regions of Central Punjab, Pakistan has reported a higher prevalence of 26.86% through PCR in bovines (n=2176) (Siddique et al., 2020). Yet another study conducted on crossbred cattle (n=100) of Sahiwal, Punjab has reported 29.0% prevalence through small subunit ribosomal RNA gene-based PCR (Chaudhry et al., 2010). A study from KP conducted on cattle (n=2,400) has reported a yet higher prevalence of 27.5% through qPCR (Pakistan, 2013). A higher percentage prevalence of 27.5% has been reported for cattle in KPK through PCR (Pakistan, 2013). A yet higher incidence of 54.8% for cattle has been reported from Baluchistan province of Pakistan (Rafique et al., 2015). Comparing our results with the work conducted in other parts of the world, it was revealed that globally higher values of prevalence have been noticed. An Iraqi study conducted on buffaloes (n=194) has reported an overall prevalence of 45.2% for babesiosis through conventional PCR (Ateaa & Alkhaled, 2019). Lower prevalence in our study attained through PCR (15.0%) as compared to previous studies could be attributed to difference in sample number, breeds, climate and types of PCR. In addition, the area under investigation (Southern Punjab) could be lesser prone to TBDs as compared to other parts of Pakistan. Lower prevalence in our study could also plausibly be attributed to better management of animals owing to better awareness of livestock farmers.

Regarding our results of prevalence for the studied cattle breeds, it was noticed that one of the animals from Cholistani breed was Babesia-positive. However, higher prevalence was noticed for crossbred cattle (46.7%) followed by that in Friesian (16.1%), Jersey (7%) and Sahiwal (6.9%) cattle breeds. Our results are in line with almost all research work published earlier. It has already been well elucidated that crossbred cattle and exotic breeds of cattle are more vulnerable to all of the TBDs including babesiosis. On the other hand, the indigenous humped zebu cattle in any part of the world (Sahiwal, Cholistani etc.) are hardy and tick-tolerant breeds hence being less prone to them (Dikmen, Dahl, Cole, Null, & Hansen, 2017; Farooq, Samad, Sher, Asim, & Khan, 2010). A work from Cholistan desert of Pakistan though conducted on theileriosis, has reported similar results (SA et al., 2016). Furthermore, the zebu cattle breeds have a potential to maintain most of their physiological (haematochemical) parameters at a harmonious pattern in all seasons, without showing much variation during stress free or stressful times (Farooq, Ijaz, Ahmed, Rehman, & Zaneb, 2012). This stress-bearing property probably renders them free from any clinical signs of TBDs as shown in our results. The exotic Taurine cattle breeds are extremely susceptible and show severe signs of the parasitism, whereas, the indicus breeds develop a strong immunity after an infection due to their innate immunity as reviewed earlier. This immunity might be another cause for absence of clinical signs in zebu cattle as seen in our results. A plausible justification based upon the principles of “Endemic Stability” to tick-borne diseases in tropics is also being presented globally. Endemic stability is an epidemiological state of equilibrium that is characterized by absence of a clinical disease instead of high incidence of infection within of set of population (Jonsson, Bock, Jorgensen, Morton, & Stear, 2012). However, a prerequisite for the achievement of this state is presence of substantial functional/innate immunity at a young age. Though no work has yet been reported on endemic stability of tick-borne diseases from Pakistan, however, the lack of clinical symptoms and low prevalence of babesiosis in our study could be attributed to this phenomenon.

Our gender-wise results indicated that females (both of cattle and buffaloes) had a higher overall prevalence of 19.5% as compared to 17.8% for males. Relationship of Babesia with gender has been studied extensively and it has been elucidated that females are more prone to babesiosis as compared to their male counterparts. Similar results have been reported from KPK with higher prevalence of babesiosis in females (41.0%) as compared to males (23.2%) (A. Khan et al., 2020). A research works from other provinces of Pakistan have also presented similar results with higher prevalence for females (Ahmad et al., 2014; Rafique et al., 2015). Gender-associated resistance or susceptibility to babesiosis has extensively been studied in lab animal models which have revealed that female Wistar rats are more susceptible as compared to males (Aguilar-Delfin, Homer, Wettstein, & Persing, 2001). Conclusions have hence been put forth that innate immunity plays a substantial role in the resistance to Babesia infection and that genetic and gender-related factors influence the efficiency of the protective response (Romero-Salas et al., 2016).

In the present study, prevalence percentage for babesiosis was ascertained as per three age groups *viz*. G1= Up to 02 Years, G2= 03-04 Years, G3= 05-06 Years. Results revealed that G1 (Up to 02 Years) showed higher prevalence of 40.0% as compared to G2 (13.1%) and G3 (8.4%), respectively. Our results are in line with most of the studies conducted on bovine population of Pakistan as well as with those conducted globally (Ahmad et al., 2014; Farooqi et al., 2017; Khattak et al., 2017). A positive correlation (r= 0.99) has been reported between TBDs and age (Lew & Jorgensen, 2005). A study conducted on bovine population of KPK and South Punjab simultaneously, reported 42.0% prevalence of TBDs in animals less than 1 year old as compared to 33.1% for those above 1 year of age (Q. U. Ashraf et al., 2013). Similarly, from KPK, a higher prevalence of babesiosis (75.3%) has been reported for young cattle elsewhere (A. Khan et al., 2020). Yet another study which incorporate three agro-ecological zones of Pakistan has reported higher prevalence in young cattle (23.1%) as compared to older cattle (11.9%) (Siddique et al., 2020).Various scientific and medical reasons have been put forth by researchers regarding this age-associate vulnerability to babesiosis and other TBDs. As per one reasoning, the young animals have thin and soft skin which is easier for ticks to infest, resulting in higher tick infestation and resultantly higher susceptibility towards TBDs including babesiosis (A. Khan et al., 2020; Zahid et al., 2005). Apart from this, failure in passive transfer (FPT) due to decreased intake of colostral antibodies by young ones may result in lesser immunity and hence a reason of susceptibility towards babesiosis. In Pakistan, the pattern of colostrum feeding is quite flawed which makes it vital to streamline efforts towards appropriate colostrum feeding and hence enhanced immunity (Lashari et al., 2020).

Summer season had highest prevalence (35.9%) of babesiosis in our study as compared to the winter season (5.7%). These results are also in concordance to earlier published work both from Pakistan and other countries of the world (Ahmad et al., 2014; Farooqi et al., 2017; Pakistan, 2013; Siddique et al., 2020). Similar to age, seasons have also been highly associated with level of prevalence of babesiosis and all other TBDs (Ahmad et al., 2014; Pakistan, 2013; Siddique et al., 2020; Zahid et al., 2005). A study from Layyah conducted on prevalence of anaplasmosis (a TBD) has reported highest prevalence during autumn (18.3%), followed by that in summer (9.7%) and winter season (7.1%) (S. Ashraf et al., 2021). Summer season allied with humidity has emerged in two summer seasons in Pakistan i.e., dry summer (May, June) and wet summer (July, August). In general, whole summer season owing to its conducive environment for growth of ticks’ results in a high tick population which ultimately results in higher infestation of livestock and higher susceptibility to TBDs including babesiosis. This has been reported from all south Asian nations such as India (Ghosh et al., 2007; Minjauw & McLeod, 2003; Singh et al., 2013). Pakistan, lying in the WCZs of the world, makes it highly susceptible to babesiosis and other TBDs.

### Hematological Profile

Amongst the studied hematological attributes, Hb concentration, TEC, PCV and MCV were lower in babesia-positive animals as compared to healthy ones. However, the TLC and lymphocytes were higher for infected animals as compared to healthy, negative ones. Our results are completely in accordance to the previously published international literature that the TBDs including babesiosis hamper significantly the blood profile of infected animals (Fadly, 2012; Javed et al., 2014; Mahmoud et al., 2015). From Pakistan however, mostly the research work conducted on TBDs is related to their prevalence and molecular identification rather than hematological assessment. A study conducted on cattle and buffalo of South Punjab regarding hematological alterations in Babesia, results similar to ours have been presented (Zulfiqar et al., 2012). Previous international reports have also clearly document a significant decrease in RBC count, Hb and PCV values in livestock with TBDs (Col & Uslu, 2007; I. A. Khan, Khan, Hussain, Riaz, & Aziz, 2011). The plausible justification for these hematological alterations in infected bovines is release of certain toxins/metabolites Babesia species., blood loss due to tick infestation, parasitemia-induced-anemia, immune-mediated erythrophagocytosis and Tumour-Necrosis Factor-α (Boulter & Hall, 1999; Geerts, Holmes, Eisler, & Diall, 2001). It has also been postulated that the Babesia species causes macrocytic hypochromic anemia, indicative of severe intravascular hemolysis of RBCs in bovines affected with persistent babesiosis (Ibrahim, El Behairy, Mahran, & Awad, 2009; Mahmoud et al., 2015; Zulfiqar et al., 2012). These may be due to the fact that although Babesia sp. may cause direct damage on some erythrocytes, immune-mediated injury of parasite may be more important in the pathogenesis of anemia (Messick, 2004). Yet, the increase in erythrophagocytosis by activated macrophages (Court, Jackson, & Lee, 2001) and the production of anti-erythrocyte antibodies (Góes, Góes, Ribeiro, & Gontijo, 2007) may also contribute to the development of anemia.

In a nutshell, the overall prevalence of babesiosis in cattle population of Southern Punjab (19.4% by PCR) is lower than other parts of Pakistan. Furthermore, this region has *Babesia bigemina* as the prevalent species. Microscopy is a less sensitive and specific test which need to be replaced with novel molecular diagnostic tests such as IFAT, ICT and ELISA. Development of an appropriate vaccine from the field strain of Babesia using proteomics and DNA technology is a need of time.

